# CodonTranslator: a conditional codon language model for codon optimization across life domains

**DOI:** 10.1101/2025.11.24.690310

**Authors:** Yanshuo Chen, Yuming Zhang, Joshua Li, Boxue Tian, Heng Huang

## Abstract

Codon optimization involves selecting synonymous codons to match host-specific preferences. It is critical for heterologous expression but remains challenging due to the combinatorial design space. Under long-term evolutionary selection, natural coding sequences are near-optimal compromises between translational efficiency, accuracy, and regulatory constraints, providing a *de facto* standard for data-driven models. Recent deep learning–based language models therefore aim to learn the distribution of natural codon sequences and reuse it for design. However, existing approaches discard the rich semantic structure of taxonomic lineages, underutilize protein functional and evolutionary constraints, and often rely on masked-language objectives that lack a principled mechanism for sequence generation. Here we present CodonTranslator, a 150M-parameter decoder-only Transformer trained on 62 million CDS–protein pairs from over 2,100 species. CodonTranslator uses a pretrained language model to embed hierarchical species lineages and a pretrained protein language model to encode protein context, enabling interpolation across hosts and generalization to unseen species and proteins. Our results show that CodonTranslator implicitly learns the genetic code from data, faithfully reproduces species-specific codon usage, and designs coding sequences that match or surpass existing methods in both codon usage metrics and predicted biological stability. Our dataset, pretrained models, and code are available at https://github.com/poseidonchan/CodonTranslator

## 1 Introduction

Synonymous codons are not used uniformly within or across genomes. This persistent heterogeneity—codon bias—arises from the interplay of mutational pressures, natural selection on translational efficiency, and genetic drift and manifests at three nested scales: (i) between species, (ii) among genes within a genome, and (iii) across sites within a single coding sequence [1,2]. At the interspecies level, compositional constraints (*e.g*., GC content), codon–context effects, and codon-pair preferences shape the accessible sequence space, generating stable, lineage-specific patterns responsive to ecological and cellular demands [3,4,5]. Within genomes, highly expressed proteins often enrich for efficiently decoded codons, reflecting selection for speed and accuracy, whereas locally non-optimal codons can be maintained when slower translation benefits cotranslational folding or assembly [4,6]. Within genes (site level), selection pressure is also uneven, since structurally or functionally sensitive regions favor codons that reduce mistranslation, while more flexible segments tolerate broader usage; accordingly, synonymous substitutions can tune ribosome traffic, folding pathways, and cellular fitness without altering the amino-acid sequence [7,8,9].

Given the impact of codon usage on protein expression, codon optimization—selecting an appropriate codon sequence for a given amino-acid sequence in a target host—is critical for synthetic biology and biotechnology. Early computational approaches relied on rule-based codon tables to guide sequence design [10,11,12,13], typically by preferring high-frequency codons in highly expressed genes. However, this design strategy has been challenged by observations that such optimized sequences can reduce protein efficacy and stability [14].

With the development of deep learning and the availability of large biological sequence databases, the field has shifted toward learning the complex “natural language” of codon usage directly from natural sequence data [15,16,17]. Several BERT-based language models have been specifically developed for codon optimization. CodonTransformer tailors a BERT model with carefully designed tokens for protein-to-codon translation across 164 species and encodes species identity via discrete embeddings [18]. Similarly, SynCodonLM employs a specialized masking scheme for the BERT model to ensure correct codon–amino acid mapping and clusters all species into 500 discrete tokens to train on over 65 million sequences [19]. Meanwhile, CodonBERT applies a cross-attention mechanism to better model protein-conditioned generation but was trained only on the human genome [20]. Together, these models improve codon optimization performance and mark a paradigm shift toward complex language modeling.

Despite recent progress, current methods still suffer from several key limitations. First, species context is poorly modeled. Most approaches represent hosts as independent categorical labels, which ignores the rich hierarchical relationships captured by taxonomy and limits generalization to unseen or under-represented species. Second, protein context is underutilized. The function and evolutionary history of a protein influence local codon choices [21,22], yet many models treat the amino-acid sequence as a simple, unconditional input. Third, the BERT-style training objective is misaligned with the sequence generation task. Theoretical analyses of masked diffusion language models show that BERT-family models, which are trained with a fixed masking ratio, are not designed to recover sequences from arbitrary mask states [23,24].

To address these limitations, we present CodonTranslator, a conditional generative language model for codon optimization. Our model is a decoder-only Transformer that generates codon sequences conditioned on both species and protein context. We represent species with continuous embeddings derived from hierarchical lineage text using the `Qwen-Embedding-0.6B` model [25], and we extract deep protein features using the `ESM-C 300M` model [26] to capture functional and evolutionary constraints. We pre-train CodonTranslator on a largescale dataset of 62 million protein–coding sequence (CDS) pairs spanning 2,163 species [27,28] to ensure robust generative performance across diverse organisms. We show that the pre-trained CodonTranslator faithfully learns natural codon usage and can generate codon sequences across all domains of life.

## 2 Results

### 2.1 CodonTranslator is a conditional codon language model

CodonTranslator formulates codon optimization as a conditional sequence generation problem: given a target host species and a protein amino-acid (AA) sequence, the model generates a host-adapted coding sequence (CDS). Analogous to neural machine translation, we adopt a prompt–response formulation in which the species and protein jointly define the prompt and the CDS is the response (Fig. 1a,d). The species prompt is constructed from hierarchical lineage text (*i.e*., kingdom → · · · → species) retrieved from GBIF (the Global Biodiversity Information Facility) (Fig. 1c), and the protein prompt is the raw AA sequence of the target protein. The model outputs a CDS as a sequence of codon-level tokens, with each three-nucleotide codon represented as a single token in the vocabulary.

**Fig 1.**
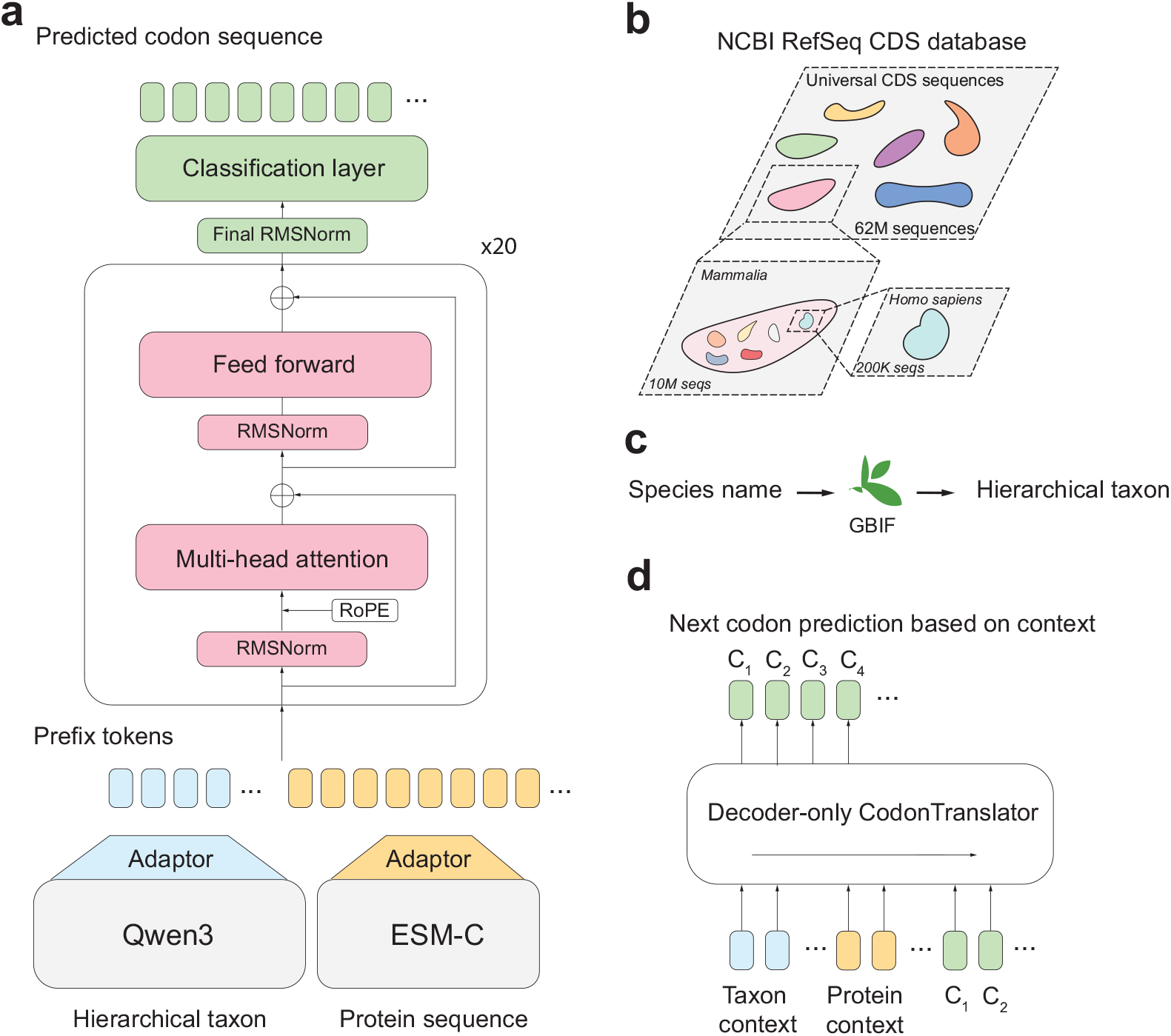
Overview of CodonTranslator architecture and training scheme. **a** Model architecture of CodonTranslator. CodonTranslator is a decoder-only transformer with two frozen encoders. The model takes a hierarchical taxon representation and a protein sequence as conditioning inputs and generates the corresponding codon sequences. **b** Training dataset landscape. The dataset includes over 62 million CDS from the NCBI RefSeq database. The hierarchical illustration reflects our hypothesis that codon usage is more similar between closely related species. **c** The hierarchical taxon representation is constructed by querying the GBIF database using the species name. **d** Next-codon prediction training scheme. Species and protein tokens are prepended to the codon sequence, and the model is trained to predict the next codon based on the preceding context.

The main architecture of CodonTranslator is a decoder-only Transformer ( ≈ 150M parameters) with two frozen pre-trained encoders (Fig. 1a) named as `Qwen3-Embedding-0.6B` [25] and `ESM-C 300M` [26]. The species and protein prompt are fed into `Qwen` and `ESM-C`, respectively, to obtain the per-token and per-AA embeddings. After that, the species and protein embeddings are independently projected to the decoder’s hidden dimension by light-weight adapter modules and serialized as prefix tokens that precede the codon tokens. The decoder itself is a pre-norm Transformer [29] consists of standard multi-head self-attention blocks [30] with rotary positional encodings [31] and a linear–softmax output head over the codon vocabulary (see more details in Methods). This conditioning scheme offers two key advantages. First, taxonomy-aware species representations enable smooth interpolation in embedding space, facilitating transfer to under-represented or entirely unseen hosts. Second, protein embeddings supply sequence-specific context beyond AA identity alone, allowing the model to tailor codon choices to the biochemical and structural demands of each protein rather than relying solely on global codon usage trends.

We trained CodonTranslator on a large-scale corpus of 62 million CDS–protein pairs spanning 2,163 species curated from the NCBI RefSeq database (Fig. 1b) [27,28]. To assess out-of-distribution generalization, we constructed an unseen-species test set by removing ten species entirely from the corpus and evaluating them only at test time. For the remaining species, data were split into training and validation sets in a 99:1 ratio at the protein level to prevent sequence leakage between splits (see more details in Methods). Taxonomic strings for all species were programmatically queried from GBIF and normalized to ensure consistent lineage construction (Fig. 1c). This setup establishes CodonTranslator as a pre-trained, taxonomy-aware language model that explicitly learns and exploits the joint dependence of codon usage on species lineage and protein context.

### 2.2 CodonTranslator learns the pattern of natural codon sequences

To evaluate whether CodonTranslator learns the patterns of natural coding sequences, we first checked its training dynamics to see whether there was any overfitting (Fig. S1a). The loss curve for CodonTranslator was stable, with monotonically decreasing validation and test losses, indicating that the model had healthy generalization ability. Based on this observation, we selected the final checkpoint for downstream analyses.

A key difference between our method and other language models is that we do not enforce that the model can only choose the correct codon during training or inference; instead, we believe that the model itself could recover the genetic code through large-scale pretraining. We then investigated whether CodonTranslator learns the genetic code by examining the next-codon probability distribution. We aggregated the next-codon probability distributions over both validation and test sequences and plotted the confusion matrix and the area under the receiver operating characteristic curve (AUROC) for each AA (Fig. S1b,c). The results show that our model can successfully distinguish codons consistent with the genetic code from those that are not: the overall AUROC is around 0.84, and the confusion matrix visually shows that the correct codon is assigned the highest probability.

Before comparing the generated codon sequences with the natural codon sequences, we first define the metrics used in Fig. 2, which quantify host-specific preferences, sequence similarity, and compositional bias. The codon similarity index (CSI) is computed as the geometric mean of species-specific codon weights along a sequence [32,33]; higher CSI indicates better alignment with the host’s preferred codons for that protein context. Importantly, a higher CSI is *not* universally optimal for expression: pushing too strongly toward highly preferred codons can increase mRNA secondary structure or disrupt beneficial ribosome pauses, so CSI needs to be interpreted together with other features [34]. To measure direct sequence recovery, codon similarity is defined as the codon-level identity fraction between the generated and ground-truth codon sequences. To capture global compositional properties, we use the effective number of codons (ENC), which ranges from 20 (strong bias) to 61 (no bias), summarizing codon diversity [35], and GC%, the nucleotide GC content of the predicted CDS, which reflects mutational biases and thermodynamic stability and interacts with codon usage. Finally, to assess local usage profiles along the gene, we compute a dynamic time warping (DTW) distance between smoothed %MinMax profiles [36] of ground truth and prediction; this measures how closely the model reproduces position-wise enrichment or depletion of synonymous choices, with lower values indicating better agreement. Together, these metrics probe global composition, local preference matching, and host-specific codon usage, providing a more complete view of biological plausibility than accuracy alone (more details are included in Methods).

**Fig 2.**
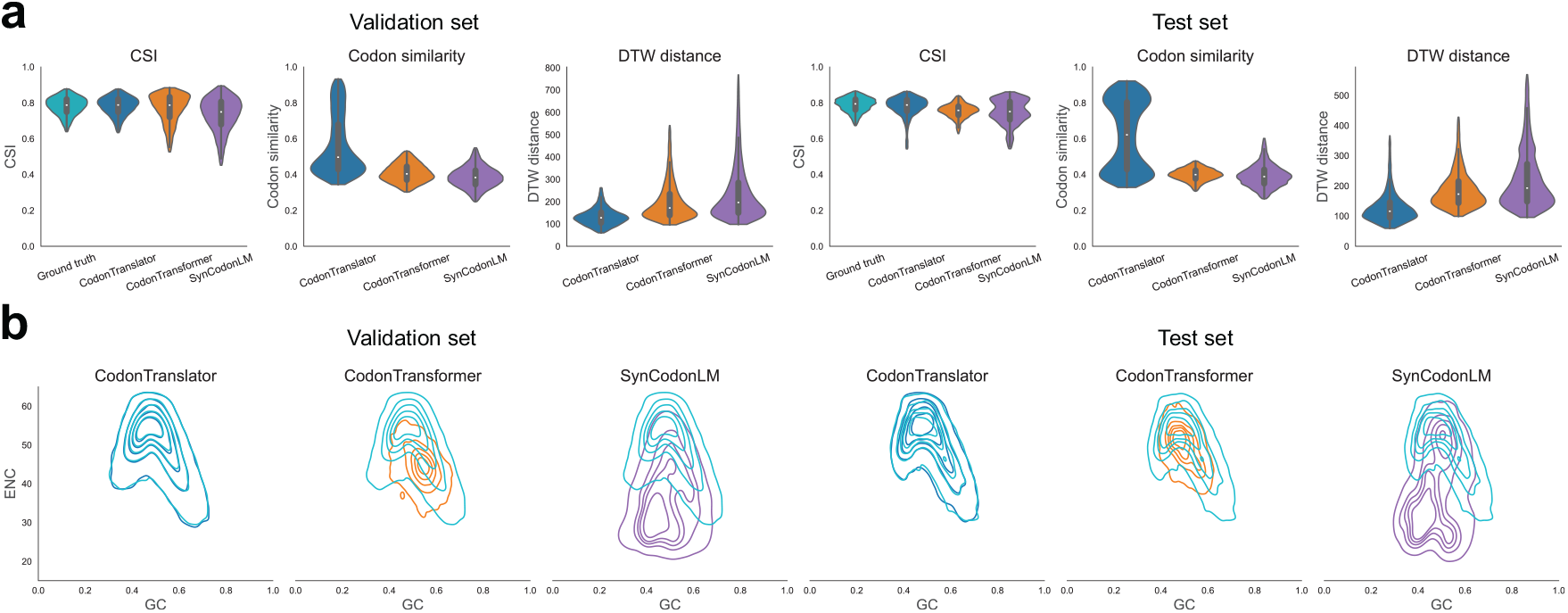
Performance of CodonTranslator on the validation and test sets. **a** CodonTranslator achieves higher codon sequence similarity and lower DTW distance compared to other models on both the validation and test sets, indicating better sequence recovery accuracy and closer resemblance to the ground truth sequences. It also outperforms other models on previously unseen species. **b** GC content and ENC landscape demonstrates that CodonTranslator accurately captures species-specific codon usage patterns, even for completely unseen organisms.

We next quantitatively assessed CodonTranslator’s performance against CodonTransformer [18] and SynCodonLM [19] on the validation and test sets using these metrics. To run these models on all 2,163 species in our dataset, we mapped each host species name to the most similar organism or cluster supported by each model and sampled sequences with temperature equals to 1. Specifically, for CodonTransformer, since it only includes 164 organisms, if no organism was close to the query species name, we assigned *Homo sapiens* as the default organism. We observe that CodonTranslator learns patterns consistent with natural sequences on both validation and test data (Fig. 2a). All three models achieve similar CSI levels, indicating broadly comparable alignment with host-specific codon preferences at the global level. However, CodonTranslator attains the highest codon similarity and the lowest DTW distance, with CodonTransformer performing slightly better than SynCodonLM on these two metrics. In the GC–ENC landscape of generated and true sequences (Fig. 2b), CodonTranslator’s outputs occupy taxon-specific regions that most closely mirror those of real coding sequences, including species that were completely unseen during training (Fig. 2b). This indicates that the taxonomy-informed species embeddings guide the decoder toward lineage-appropriate composition and bias, rather than a single universal solution. Together, these results support the view that CodonTranslator captures the multi-scale structure of codon usage—from overall base composition to local codon-pair and position-wise preferences—and that its conditioning scheme enables meaningful generalization to novel hosts without relying on hand-crafted codon vocabularies or species-specific rules.

### 2.3 Model benchmark on model organisms

We then compare CodonTranslator with CodonTransformer, SynCodonLM, and three widely used vendor algorithms—Twist Bioscience, Integrated DNA Technologies (IDT), and Genewiz—focusing on five canonical model organisms: *Escherichia coli, Saccharomyces cerevisiae, Arabidopsis thaliana, Mus musculus*, and *Homo sapiens* (Fig. 3). This setting mirrors practical codon optimization workflows, where traditional tools are available primarily for model hosts. To ensure a fair comparison, we evaluate all methods on the same protein sets used in the CodonTransformer paper [18] and report three complementary metrics: codon similarity, DTW distance between smoothed %MinMax usage profiles, and CSI. In addition, the GC% and ENC landscape is examined in Fig. S2 to assess whether a method reproduces each organism’s complex codon-usage pattern. Together, these readouts probe exact recovery (codon similarity), local usage trajectories (DTW), and global compositional and bias properties (CSI, GC, and ENC), providing a balanced picture of biological plausibility rather than a single scalar score.

**Fig 3.**
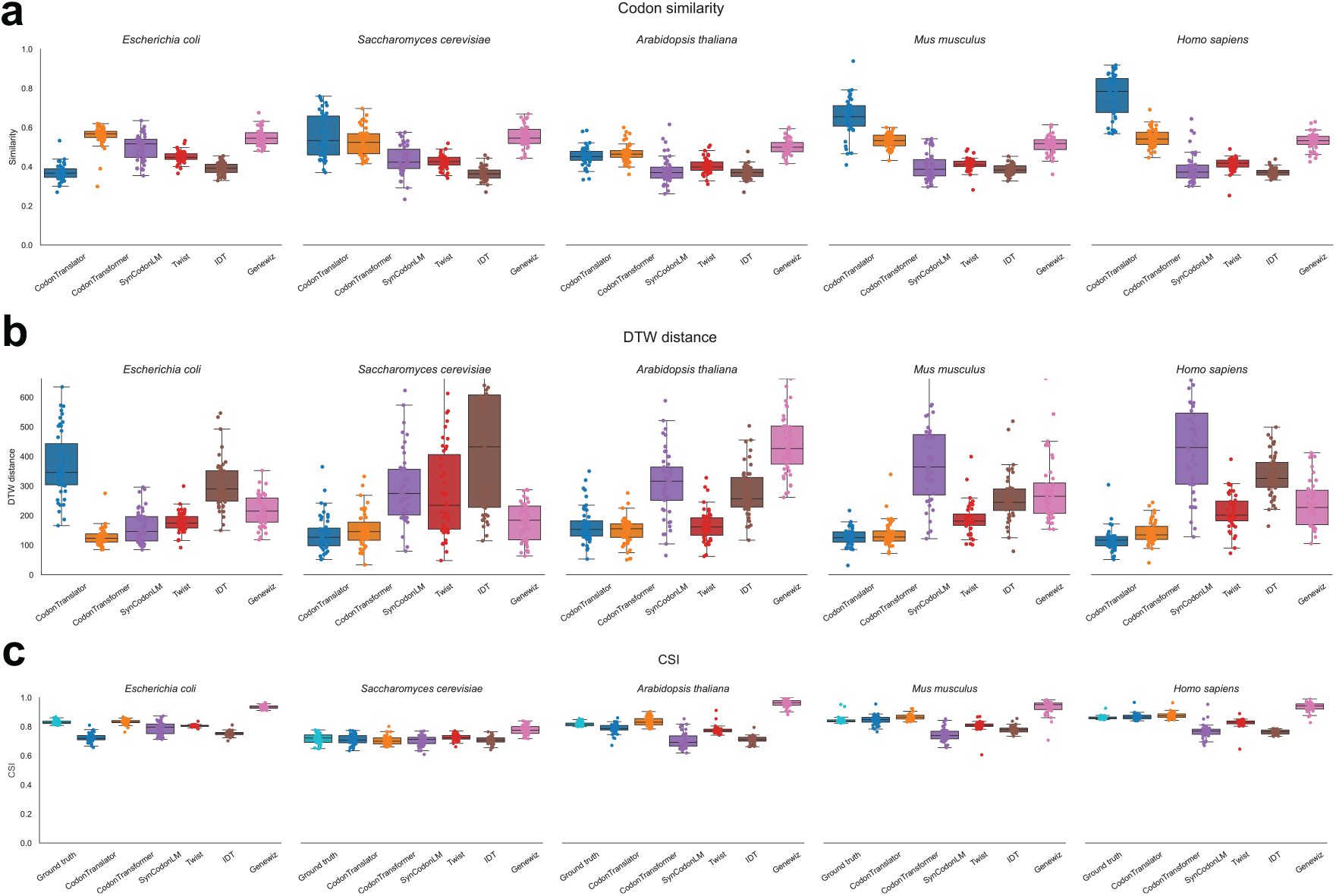
Benchmarking of codon sequence optimization methods on model organisms. Performance of six codon optimization methods on five model organisms. **a**, codon similarity (fraction of codons identical to the native CDS); **b**, DTW distance between smoothed %MinMax usage profiles; and **c**, Codon Similarity Index (CSI). Box plots summarize the distribution across 50 genes for each method: the central line denotes the median, the box spans the interquartile range (25th–75th percentile), and overlaid points represent individual genes.

Across organisms, CodonTranslator achieves high codon similarity and is particularly strong on the mammalian hosts *Homo sapiens* and *Mus musculus*, where the interquartile ranges are visibly elevated relative to the other methods (Fig. 3a). For *Saccharomyces cerevisiae* and *Arabidopsis thaliana*, CodonTranslator remains competitive, with median similarities comparable to or higher than those of CodonTransformer and SynCodonLM. On *E. coli*, vendor algorithms and CodonTransformer reach higher codon similarity, but CodonTranslator still maintains reasonable identity to the native sequences. Local usage trajectories show a similar pattern. CodonTranslator attains low DTW distances to the ground truth across most organisms (Fig. 3b), indicating that it can reproduce the spatial pattern of enrichment and depletion of synonymous codons along a gene. This profile-level agreement matters in practice: ribosome traffic, co-translational folding, and mRNA surveillance respond to position-dependent usage rather than only global composition. For the four non-bacterial hosts, CodonTranslator generally matches or improves upon the learning-based baselines and vendor tools. For *E. coli*, however, CodonTransformer achieves the best performance. Global composition and bias patterns are broadly consistent with these observations. For CSI (Fig. 3c), all methods reach similar organism-specific ranges, but CodonTranslator tends to stay close to the distributions observed for native coding sequences rather than indiscriminately maximizing the index. The ENC–GC landscape comparison (Fig. S2) shows that CodonTranslator’s contours overlap the ground-truth contours closely. This analysis also reveals that vendor codon optimization methods tend to either uniformly choose synonymous codons (Twist) or strongly favor a small subset of codons (Genewiz), which likely explains why Genewiz attains very high CSI scores. Overall, the BERT methods align the ENC–GC landscape more faithfully than the traditional codon optimization tools provided by DNA synthesis vendors. These results suggest that CodonTranslator is competitive with established tools, performs particularly well on mammalian hosts, and reconstructs organism-appropriate local usage patterns and global compositional bias.

### 2.4 CodonTranslator leverages species context and generates stable sequences for heterologous expression

We first asked whether the species representations used to condition CodonTranslator capture meaningful lineage structure and are actually used by the decoder. To visualize the species space encoded by `Qwen3-Embedding-0.6B`, we extracted and averaged the species tokens for each organism into a single vector and projected these embeddings to two dimensions using UMAP (Fig. 4a, Fig. S4a; *n* neighbors = 15, min dist = 0.5). Coloring by kingdom and phylum yields coherent clusters that align with major taxonomic divisions, suggesting that the hierarchical lineage text preserves phylogenetic proximity. At higher resolution (Fig. S4a), class and order remain separable, and a circular phylogenetic rendering of all 2,163 species (Fig. S4b) mirrors the embedding neighborhoods. These observations indicate that continuous species embeddings respect parent–child relationships rather than collapsing host identity to isolated one-hot labels, and they provide a natural inductive bias for interpolating to under-represented or unseen hosts.

**Fig 4.**
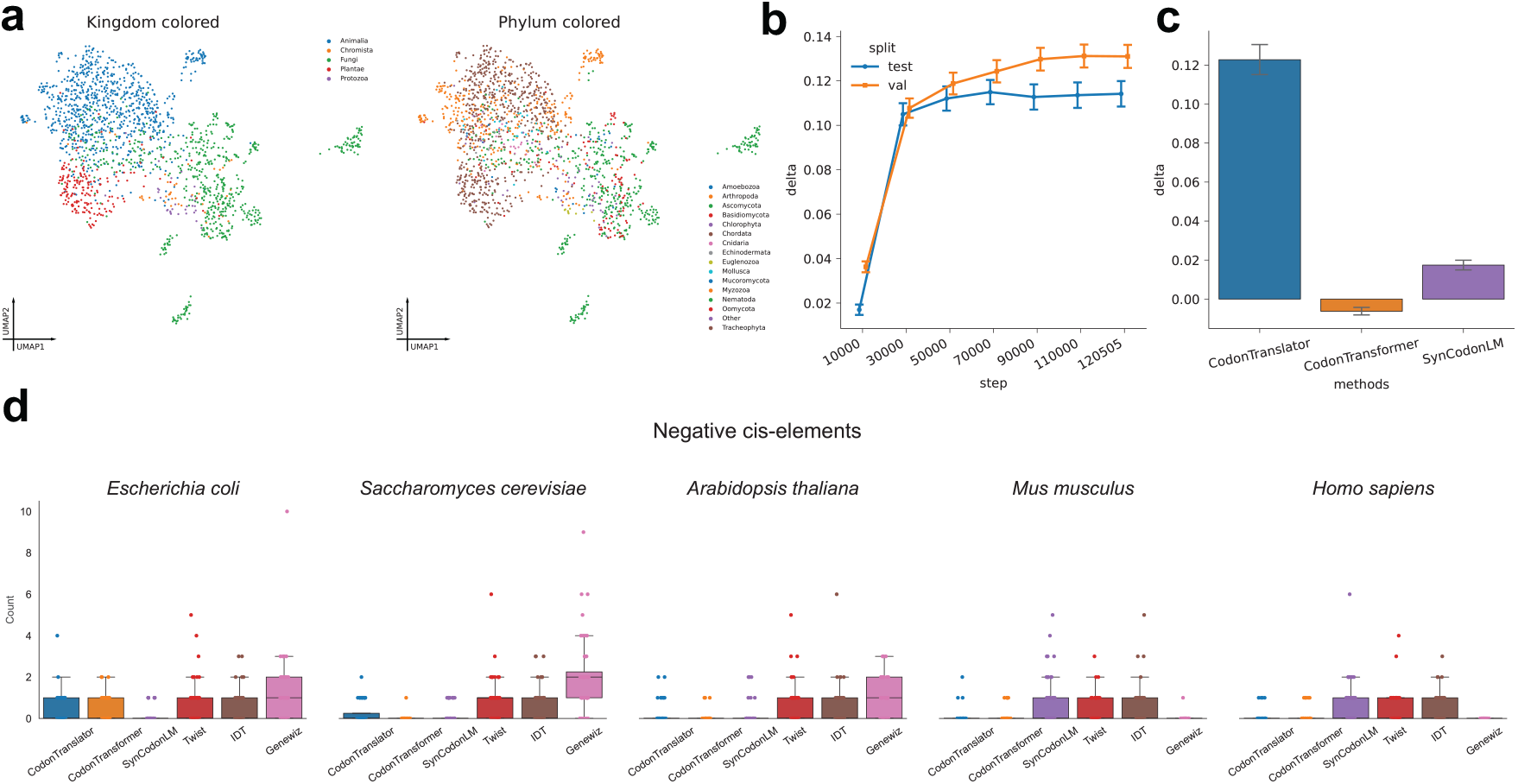
CodonTranslator effectively captures species-specific information and enhances heterologous expression. **a** UMAP visualization of species embeddings colored by Kingdom and Phylum, revealing clustering based on taxonomic relationships. **b** Effect of species information ablation on performance at different training checkpoints, quantified by the change in codon seqeunce similarity (Δ) when using correct species information. A sharp delta increment around 20,000 steps indicates the model begins to learn the species condition at early stage. **c** Comparative ablation study across models. The pronounced performance gain in CodonTranslator when using corresponding species underscores its strong reliance on species information, while this dependency is less evident in CodonTransformer and SynCodonLM. **d** Evaluation of negative *cis*-elements in 52 proteins commonly used for heterologous expression. A higher score indicates lower predicted expressibility. Box plots summarize the distribution across 52 genes for each method: the central line denotes the median, the box spans the interquartile range (25th–75th percentile), and overlaid points represent individual genes.

To test whether the decoder actually uses this species information during learning, we designed a species ablation protocol and tracked its effect across training checkpoints (Fig. 4b). From the validation and test sets, we sampled roughly 2,000 proteins and, at each checkpoint, generated two codon sequences per protein: a conditional sequence using the correct species prompt, and an ablated sequence in which the species prompt was replaced by a random species drawn 10 times. We then computed the change in codon similarity Δ (original species minus ablated), so larger positive values indicate stronger reliance on the species prompt. Early in training, Δ is near zero, consistent with the decoder initially fitting generic nucleotide statistics. After 10,000 steps, Δ first increases sharply and then gradually increases, showing that the model progressively learns to exploit species-specific cues while the protein prompt remains fixed. This trajectory provides a causal signal that lineage information becomes operational in the generative pathway, rather than acting as a passive tag.

We also compared species sensitivity across modeling paradigms by applying the same ablation protocol to CodonTransformer and SynCodonLM (Fig. 4c). CodonTranslator shows the largest drop in performance when the species prompt is randomized, indicating that its outputs depend strongly on the supplied host. CodonTransformer exhibits a weaker response, consistent with its use of a discrete host token and its practical fallback of mapping unfamiliar species names to a small set of common organisms before generation, which compresses species variation and makes ablation less disruptive. SynCodonLM shows an intermediate pattern, as clustering all species into a limited codebook improves robustness but reduces organism-level resolution. Together with the manifold analyses above, these results support the idea that continuous, taxonomy-aware conditioning is important for faithfully expressing inter-species differences in codon usage.

In many practical scenarios, however, the goal is heterologous expression in a model host rather than recovery of the native sequence. In such cases the species prompt will deliberately not match the natural organism, and the question becomes whether the generated CDS is biologically reasonable for the chosen host. To probe this setting, we carried out an *in silico* benchmark on 52 widely used recombinant proteins expressed in five model organisms and quantified predicted negative *cis* elements in the CDSs generated by each method using GenRCA [37] (Fig. 4d). On this measure, CodonTranslator and CodonTransformer reduce the *cis*-element burden compared to DNA synthesis vendor tools, with the largest gains in human, mouse, and yeast, while SynCodonLM remains strongest in *E. coli*. Thus, even when the conditioning species is intentionally changed (*e.g*. heterologous design scenario), CodonTranslator produces sequences with fewer predicted liabilities in eukaryotic hosts.

## 3 Discussions

In this work, we introduced CodonTranslator, a conditional codon language model that generates host-adapted coding sequences given a species and a protein sequence. The model combines frozen, pre-trained species and protein encoders with a decoder-only Transformer trained on 62 million CDS–protein pairs across 2,163 species. We showed that CodonTranslator learns the genetic code directly from data, recovers species-specific codon usage on validation and held-out species, and is on par with or surpasses existing Transformer-based models and traditional vendor tools across diverse metrics. We further demonstrated that the model can be used in heterologous expression tasks to generate biologically stable coding sequences.

A key contribution of this work is the use of continuous, taxonomy-aware species embeddings as conditioning signals for codon optimization. By encoding hierarchical lineage strings with a large language model, CodonTranslator places species in a smooth embedding space where taxonomic neighbours cluster together. Species ablation experiments across training checkpoints further validate that the model progressively learns to use this information. Another advantage is the use of a decoder-only Transformer instead of previous encoder-only models [18,20,17,19]. This shift is important for generative codon design, since BERT-like models cannot natively sample sequences at inference time [23,24]. Our experiments show that CodonTranslator captures natural codon distributions more faithfully than BERT-based models (Fig. 1b).

Although CodonTranslator is explicitly trained on species–protein–CDS triplets, we also find that it generalizes beyond this setting. On species that are completely unseen during training, the model produces sequences that fall into the correct GC–ENC regions and match native codon usage at both global and local levels. In heterologous expression benchmarks, CodonTranslator and CodonTransformer reduce predicted negative *cis* elements compared with vendor tools in human, mouse, and yeast, while maintaining native-like stability profiles.

In conclusion, CodonTranslator provides a practical framework for integrating continuous species information together with deep protein feature for scalabaly designing host-adapted coding sequences. Since it learns species-specific codon usage directly from large-scale data and generalizes to unseen and heterologous hosts, we believe CodonTranslator can become a useful backbone for future codon design pipelines and broader sequence engineering tasks that require flexible, host-aware generation.

## 4 Methods

### 4.1 Problem formulation

Since there is no gold-standard definition of an “optimized” codon sequence [14], we adopt the working hypothesis that natural coding sequences are near-optimal outcomes of long-term evolutionary selection. Under this hypothesis, we cast codon optimization as a conditional codon sequence generation problem. Each training sample consists of a species label *s*, a protein amino-acid (AA) sequence *a*_1:*L*_, and a coding DNA sequence *d*_1:*N*_, where *N* = *L* + 1 when codons are counted including the stop codon, and translating *d*_1:*N*_ with the standard genetic code yields *a*_1:*L*_. The model aims to learn a left-to-right distribution over codons conditioned on the protein and species:

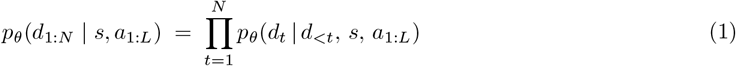

After fitting this distribution to natural sequences, we can generate candidate codon sequences for a given target host species and protein by sampling from *p*_*θ*_(*d*_1:*N*_ | *s, a*_1:*L*_).

### 4.2 Dataset construction

We curated a large-scale corpus of CDS paired with their corresponding proteins from NCBI RefSeq [27,28]. For each RefSeq record, we extracted the taxon label, the CDS nucleotide sequence, and the translated amino-acid sequence, and wrote them to parquet shards to facilitate out-of-core processing. We removed entries with empty or non-canonical coding material, CDS lengths not divisible by three, and sequences with an internal standard stop codon. To ensure robust estimation of organism-specific codon statistics, we filtered out species containing fewer than 1,000 CDS and retained a final set of 2,163 species with a total of *>*62 million CDS–protein pairs. Taxonomic strings were normalized to produce a deterministic species key used throughout preprocessing and splitting.

To evaluate generalization to unseen hosts, we defined a species held-out set (*test set*) comprising ten taxa spanning mammals, amphibians, fungi, and algae: *Ursus maritimus, Chrysochloris asiatica, Pteropus vampyrus, Xenopus tropicalis, Actinia tenebrosa, Aspergillus glaucus, Thyridium curvatum, Exserohilum turcica, Penicillium arizonense*, and *Galdieria sulphuraria*. All records from these species were excluded from training and reserved for evaluation only. From the remaining species, we constructed two disjoint subsets by selecting 1% sample for the validation set and use the 99% for training.

### 4.3 CodonTranslator framework

#### Model architecture

CodonTranslator is a decoder-only Transformer conditioned on species and protein context (Fig. 1a). It uses two pre-trained encoders, Qwen3-Embedding and ESM-C, to obtain species and protein representations, which are then passed through separate lightweight adapters. The backbone is a 20-layer Transformer with 15 attention heads and a hidden size of 750. Each layer follows a pre-norm design [29] with RMSNorm [38], multi-head self-attention [30] using rotary position embeddings (RoPE) [31] on queries and keys, and a SwiGLU [39] feed-forward network with an MLP expansion ratio of 3.2. A final RMSNorm feeds into a linear classification head that maps hidden states at codon positions to the codon vocabulary. Our preliminary experiments indicated that, for a fixed parameter budget, increasing depth improves how quickly and accurately the model learns the genetic code. We therefore adopt a configuration that is deeper than BERT-base but slightly narrower in width.

#### Model input

To encode species context, we first construct a compact textual description for each species by querying the GBIF database using the scientific species name. The description includes the full hierarchical taxonomic lineage (*i.e*. kingdom–species path) information in text. Then this text is used as the query in an instruction-style prompt:

~~~
Instruct: Given a species taxonomy information, generate a biological embedding representing its taxonomic and evolutionary characteristics.
Query:     [taxonomy description].
~~~

The instruction plus query string is embedded with `Qwen3-Embedding-0.6B`, which produces per-token hidden states 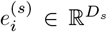, where *D*_*s*_ is the `Qwen` embedding dimension. Protein context is provided by a frozen 300M `ESM-C` encoder: given an AA sequence *a*_1:*L*_, `ESM-C` encodes it and returns per-residue embeddings 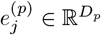 after removing special tokens, where *D*_*p*_ is the `ESM-C` embedding dimension.

Both species and protein embeddings are mapped into the Transformer hidden space of dimension *H* (here *H* = 750) by small modality-specific adapters of the same form:

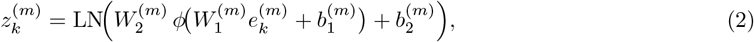

where *m* ∈ {species, protein}, *ϕ* is the ReLU activation function, 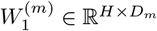 projects from the encoder dimension *D*_*m*_ ∈ {*D*_*s*_, *D*_*p*_} to *H*, and 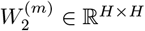 is a small bottleneck layer. The `Qwen3-Embedding` and `ESM-C` parameters are frozen; only the adapter parameters 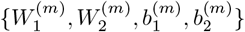 are trained.

In practice, both species and protein prefixes have variable length. Within each mini-batch, we pad the corresponding token sequences along the time axis to the batch-wise maximum, using all-zero vectors for padded positions. After passing through the adapters, these padded rows remain zero and are explicitly treated as padding. When constructing the prefix for a given example, we discard any trailing all-zero rows before concatenating the start embedding and codon tokens. As a result, padding never carries semantic signal and effectively acts as a neutral “null” context during both training and sampling.

To keep memory usage and attention cost bounded, we impose simple length caps. Protein prefixes are truncated to at most 1,024 residues before ESM-C encoding and projection. After concatenating species tokens, protein tokens, the start embedding, and codon tokens, we ensure that the total context length for each example does not exceed 2,048 positions. If the combined species+protein prefix would exceed this budget, we truncate the prefix first; if the codon sequence does not fit in the remaining capacity, we then truncate the codon side accordingly. All training examples therefore lie within the same 2,048-token context window used by the backbone.

#### Model training and sampling

We use a minimal codon tokenizer with a vocabulary of 68 tokens: four special symbols (PAD, UNK, BOS, EOS) and the 64 DNA 3-mers over { *A, C, G, T* }. Coding DNA sequences whose length is divisible by three are segmented into codons and mapped to token IDs, and an EOS token is appended by the data pipeline. At run time we do not prepend BOS; the learned start embedding inserted between the conditioning prefix and the first codon marks the beginning of the decoded sequence. The model is trained with teacher forcing to minimize the negative log-likelihood in Eq. (1), i.e. the cross-entropy between the predicted next-codon distribution and the ground-truth codon at each position. Loss is computed only on valid codon positions, ignoring PAD and EOS tokens. Species and protein encoders are frozen, while the adapter layers, codon embedding table, Transformer backbone, and output projection are updated. The length caps described above (protein prefix ≤ 1,024 residues and total context ≤ 2,048 tokens) are enforced throughout training.

We train for 3 epochs over the corpus with an effective global batch size of 1,536 sequences, obtained by combining per-device batch size, multi-GPU data parallelism, and gradient accumulation. This schedule corresponds to 120,505 optimizer steps. The learning rate is linearly warmed up from 0 to 7 × 10^−5^ over the first 10% of steps and then linearly decayed back to zero over the remaining steps. Optimization uses AdamW [40] with zero weight decay and (*β*_1_, *β*_2_) = (0.9, 0.95), together with gradient norm clipping at 1.0 to prevent rare exploding updates. Training is performed in bfloat16 mixed precision using PyTorch Fully Sharded Data Parallel (FSDP) [41] for parameter sharding, and we rely on FlashAttention’s scaled dotproduct attention kernels for efficient self-attention over long contexts. Under this setup, training the main CodonTranslator model requires approximately 2,000 NVIDIA H100 GPU-hours.

At inference time, we generate codon sequences by autoregressively sampling from the learned distribution *p*_*θ*_(*d*_*t*_ | *d*_*<t*_, *s, a*_1:*L*_). Unless otherwise noted, we use a sampling temperature of 1.0 for CodonTranslator and apply standard top-*k* and top-*p* filtering to control diversity. To ensure that the generated DNA remains consistent with the target protein, we explicitly enforce the genetic code at every decoding step: for the current amino acid *a*_*t*_, we identify the set of codons that translate to *a*_*t*_ under the standard genetic code and set the logits of all other codons to −∞ before applying the softmax, so that only genetically valid codons can be sampled at that position. The backbone uses RoPE [31] for all tokens, so positional information is encoded in a relative, multiplicative form rather than via learned absolute position embeddings. Training examples are capped at a total context length of 2,048 tokens, but at sampling time we do not explicitly enforce this cutoff and allow decoding to proceed until a designated EOS token is generated or a user-specified maximum codon length is reached. In principle, RoPE can extrapolate to contexts longer than those seen during training, and our implementation can generate sequences beyond the 2,048-token training window, although we do not systematically characterize performance in this extreme regime.

### 4.4 Evaluation metrics

#### CSI

We quantify host preference alignment with a CSI-style index defined as the geometric mean of codon weights along a CDS [32,33]. Let *c*_1_, …, *c*_*L*_ be the in-frame sense codons (stops removed). For host *h*, assign weights *w*_*h*_(*c*) ∈ (0, 1] normalized within each synonymous family (the most preferred codon has weight 1). The index calculated as:

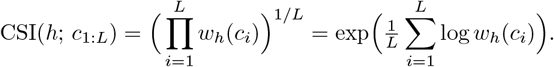

#### Effective number of codons (ENC) and GC landscape

Global codon-usage bias is summarized by Wright’s effective number of codons (*N*_*c*_, “ENC”), which ranges from 20 (one codon per amino acid) to 61 (no bias). For a synonymous family with *n* codons and counts 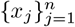 (excluding ATG, TGG, and stops), let 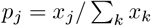 and

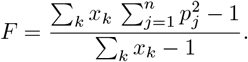

Aggregating over 2-, 3-, 4-, and 6-fold families yields

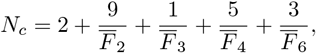

clipped to [20, 61] [35]. In parallel, we compute GC fraction as GC% = (#G+#C)*/L* over the CDS. For each method and organism we visualize the joint ENC–GC landscape and compare it with the native distribution.

#### %MinMax profiles and DTW distance

To compare position-wise synonymous usage, we use %MinMax profiles [36]. For host *h* and amino acid *a*, we estimate synonymous frequencies *f*_*h*_(*c* | *a*) from native CDS and define *f*_min_, *f*_max_, and *f*_avg_ as the minimum, maximum, and mean across codons for *a*. For a site using codon *c*,

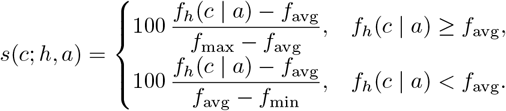

We smooth *s* along the CDS with a sliding window (default *w* = 18 codons) to obtain a %MinMax profile **p** ∈ ℝ^*L*−*w*+1^. Agreement between prediction and ground truth is measured by Dynamic Time Warping (DTW) distance between their profiles, using a squared-error local cost and monotone warping.

#### Codon similarity

Exact sequence recovery is measured as codon-level identity. Let **c**^(gt)^ and **c**^(pred)^ be the ground-truth and predicted codon sequences. We align them and let *L* be the aligned length in codons. Codon similarity is

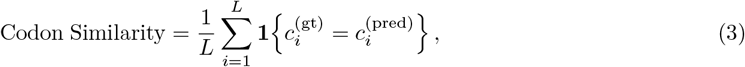

*i.e*., the fraction of positions with identical codons. This metric captures exact agreement in synonymous choices given a fixed amino-acid sequence and complements amino-acid–level and profile-level measures.

## 5 Acknowledgements

## 6 Author contributions

Y.C., Y.Z., and H.H. conceived the idea of this project. Y.Z. cleaned and curated the data. Y.C. built the model and the codebase. Y.C. and Y.Z. conducted the experiments and interpreted results. Y.C. and Y.Z. wrote the manuscript. B.T. provides helpful comments. H.H supported the research. All authors reviewed the manuscript.

## Notes

### Competing Interest Statement

The authors have declared no competing interest.

