## Supplementary information for "CodonTranslator: a conditional codon language model for codon optimization across life domains"

Yanshuo Chen[1,2], Yuming Zhang[3,4], Joshua Li[5], Boxue Tian[3], and Heng Huang[1,2,6]
[1] Department of Computer Science, University of Maryland, College Park, MD, 20742, USA.
[2] Center for Bioinformatics and Computational Biology, College Park, MD, 20740, USA.
[3] MOE Key Laboratory of Bioinformatics, State Key Laboratory of Molecular Oncology, School of Pharmaceutical Sciences, Tsinghua University, Beijing, 100084, China
[4] Peking University–Tsinghua University–National Institute of Biological Sciences (PTN) Joint Graduate Program, School of Life Sciences, Tsinghua University, Beijing, 102206, China
[5] Thomas Jefferson High School for Science and Technology, Alexandria, VA, 22312, USA
[6] Institute for Health Computing, University of Maryland, North Bethesda, MD, 20852, USA

#### Table of contents

**Supplementary Figures....................................................................................................2**

### Supplementary Figures

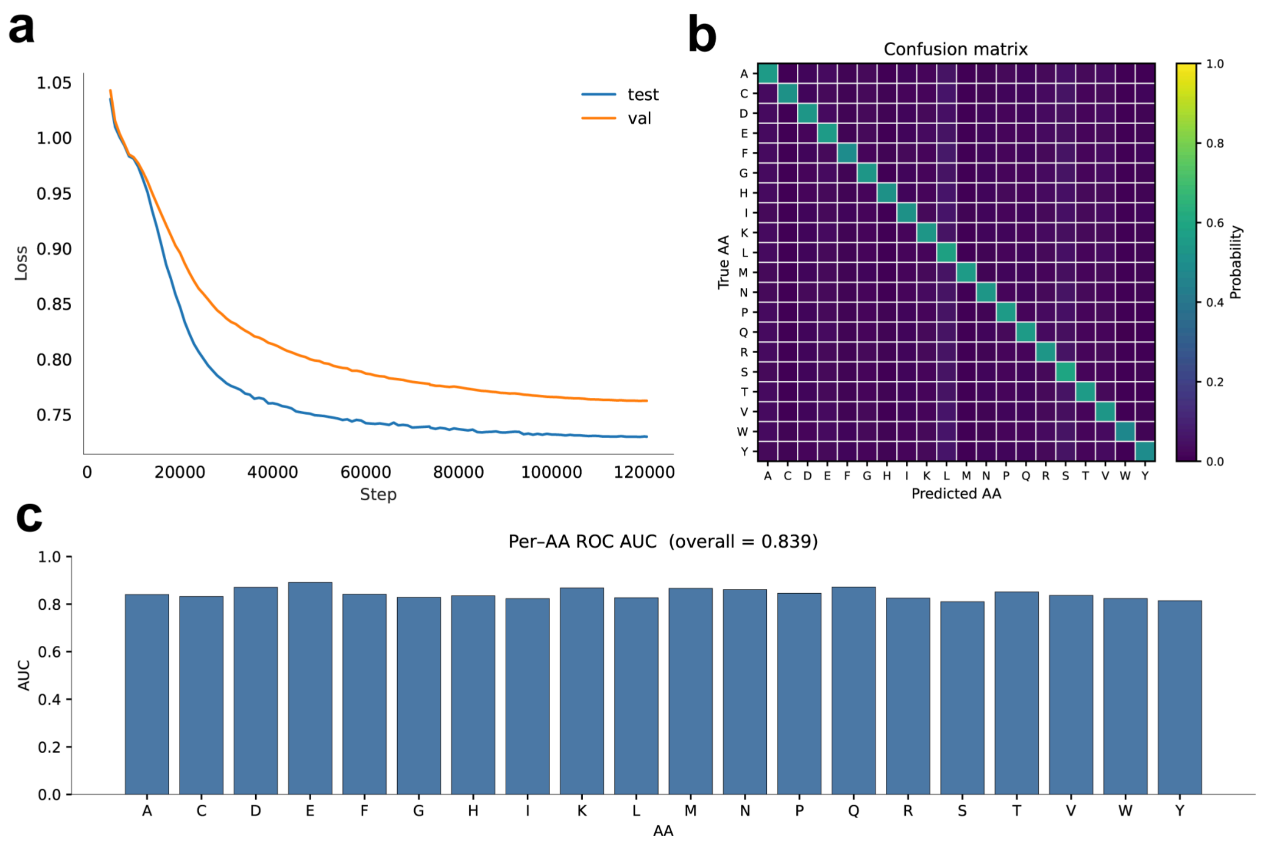


**Fig. S1. CodonTranslator training dynamics and residue-level fidelity.**
(a) Train/validation loss versus optimization steps. Loss decreases steadily with a small late-stage generalization gap, indicating stable training. The training process required approximately 20,000 GPU-hours on a cluster of NVIDIA H100 GPUs.
(b) Confusion matrix across the 20 amino acids, computed by mapping the model's predicted codons back to amino-acid classes at each position of the validation proteins. The strong diagonal and minimal off-diagonal mass show that CodonTranslator—conditioned on species and protein sequence—rarely proposes codons encoding the wrong residue.
(c) Per-amino-acid ROC-AUC; the overall AUC is 0.839, the best among all models we evaluated (including SynCodonLM and CodonTransformer), with uniformly high AUCs across residues. Guided by these diagnostics and our checkpoint sweep, we use a late-training checkpoint (~110–120k steps) for all downstream analyses reported in the main text.

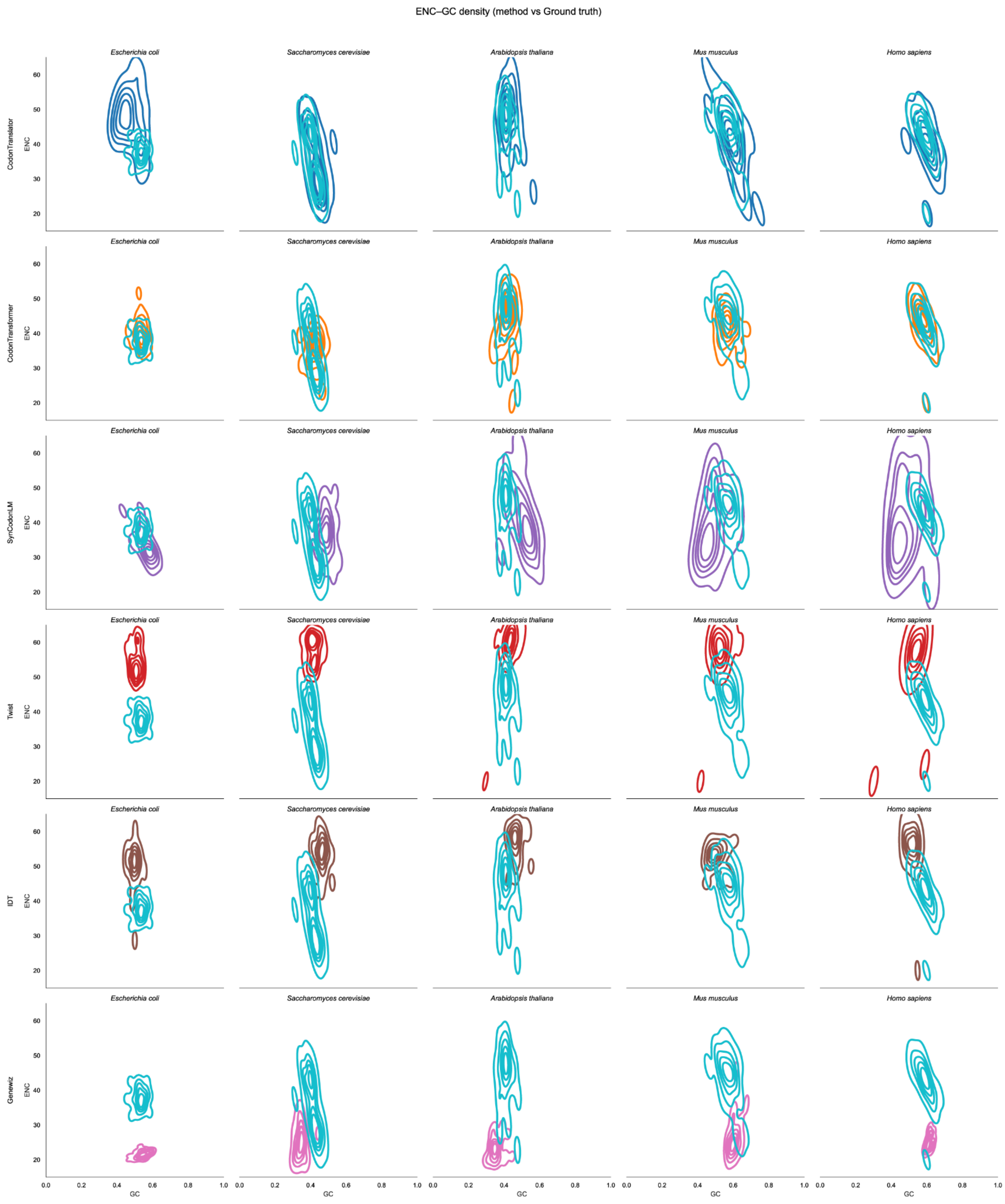


**Fig. S2. Joint ENC–GC distributions of predicted vs. native CDS across organisms.** Panels show 2-D kernel-density contours of **ENC** (Wright's Nc; y-axis, 20–61; lower = stronger codon bias) against **GC fraction** (x-axis, 0–1) for five model organisms (columns). Each row corresponds to one method and the bottom row shows the **ground truth (GT)** distributions.

Greater overlap between a method’s contours and GT indicates better recovery of the native codon-usage manifold. As expected, GC—positively associated with mRNA structural stability—and ENC—capturing codon diversity/bias—jointly delineate organism-specific optima: extremely high GC (very stable RNAs) or very low ENC (over-biased usage) need not be optimal. Across species, CodonTranslator’s organism-conditioned decoding most closely matches the GT ENC–GC landscape, whereas organism-neutral variants tend to drift toward higher ENC (weaker bias).

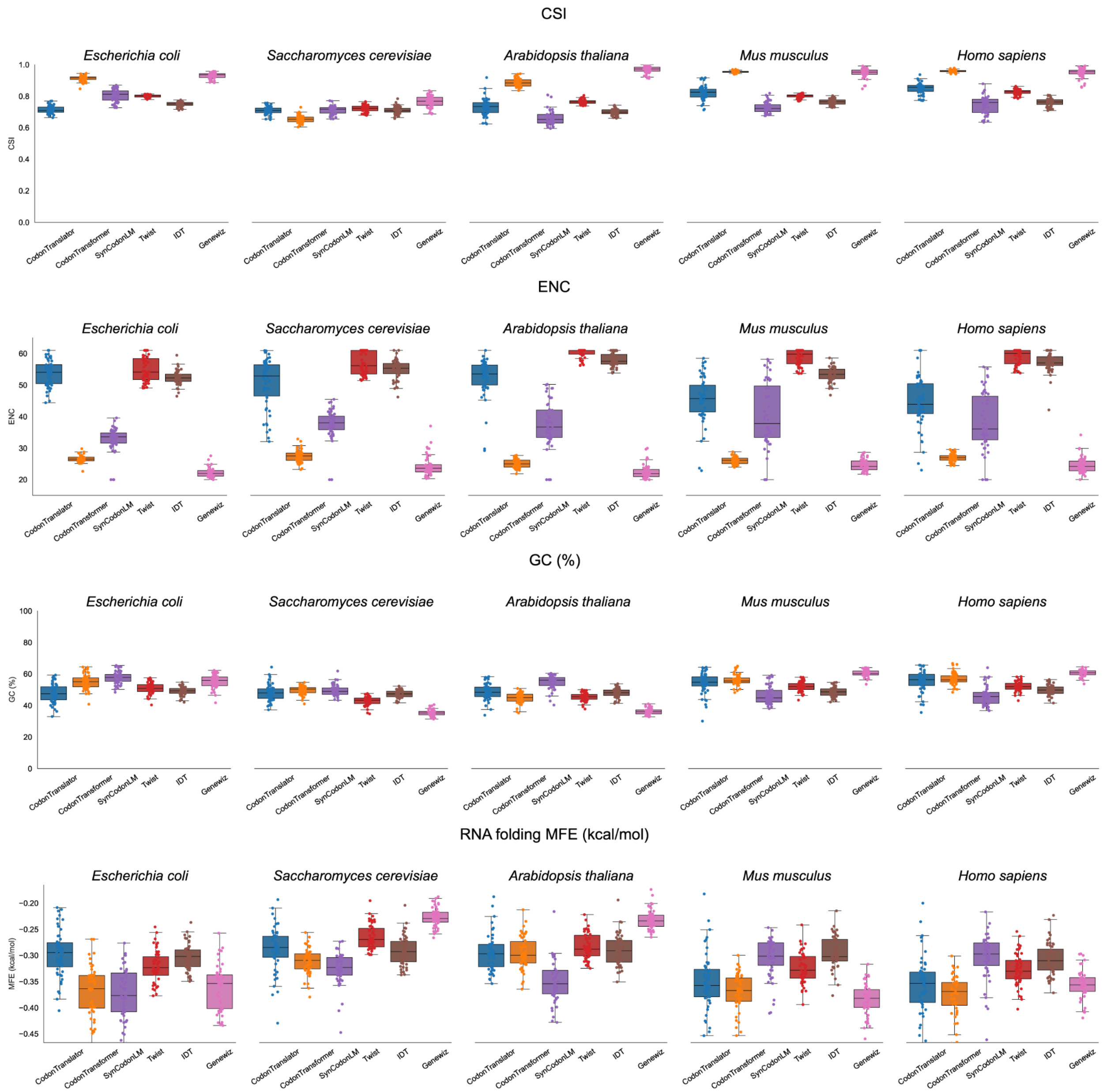


**Fig. S3. Performance of codon optimization models across five model organisms.** Boxplots summarize per-gene distributions for Escherichia coli, Saccharomyces cerevisiae, Arabidopsis thaliana, Mus musculus, and Homo sapiens comparing CodonTranslator, CodonTransformer, SynCodonLM, Twist, IDT, Genewiz and ground truth sequences on four metrics: CSI, computed as the geometric mean of species-specific codon weights—higher indicates closer alignment to host preference; ENC; GC% of the CDS; and RNA folding MFE evaluated with ViennaRNA at 37 °C and normalized by length (kcal/mol/nt). Points show individual genes; center lines indicate medians. Across species, CodonTranslator typically attains CSI comparable to or exceeding ground truth while keeping GC% near ground truth and maintaining ENC within a non-collapsed range; normalized MFE values remain within the ground truth bandwidth, indicating no reliance on excessively stabilized mRNA structure.

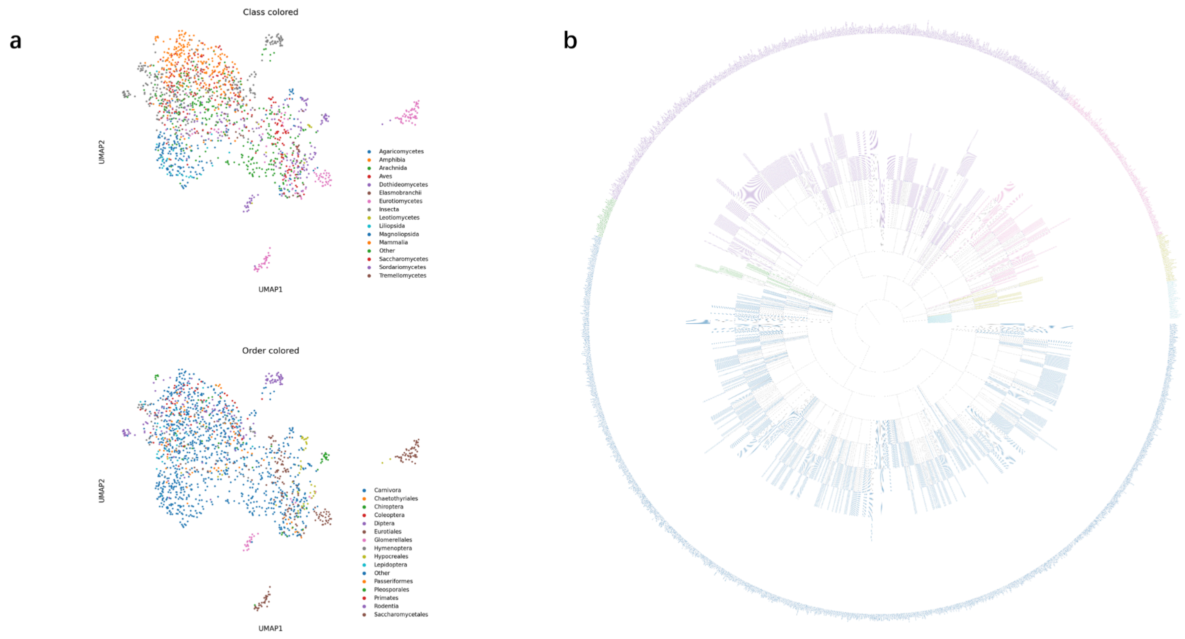


**Fig. S4. Taxonomy-aware structure of the dataset.**
(a) UMAP of species embeddings colored by Class and Order. Clear clusters align with major clades (e.g., Mammalia, Insecta, Saccharomycetes), showing that the learned species representations preserve coarse and mid-level phylogeny. We omit lower ranks (Family/Genus/Species) in panel a because the number of categories is very large.
(b) Circular phylogenetic tree for all 2,163 species in our benchmark, rendered with a Python phylogeny toolkit from NCBI taxonomy parent–child relationships; colors match panel a. A full-resolution version of the tree is provided in our GitHub repository.
